# Rational design of a novel pollinator interaction

**DOI:** 10.1101/123604

**Authors:** Kelsey J.R.P. Byers, H.D. Bradshaw

**Affiliations:** Department of Biology, University of Washington, Seattle, WA, 98195, USA

**Author notes:** Present address: Department of Zoology, University of Cambridge, Downing Street, Cambridge, CB2 3EJ, UK.

## Abstract

Diversification of the *ca*. 250,000 extant flowering plant species has been driven in large part by coevolution with animal pollinators. A recurring pattern of pollinator shifts from hummingbird to hawkmoth pollination has characterized plant speciation in many western North American plant taxa, but in the genus *Mimulus* (monkeyflowers) section *Erythranthe* the evolution of hawkmoth pollination from hummingbird-pollinated ancestors has not occurred. We manipulated two flower color loci and tested the attractiveness of the resulting four color phenotypes (red, yellow, pink, white) to naïve hawkmoths. Hawkmoths strongly prefer derived colors (yellow, pink, white) over the ancestral red, and prefer the two-locus change (white) to either of the single-locus changes (yellow, pink). The simple flower color genetics underlying this innate pollinator preference suggests a potential path for speciation into an unfilled hawkmoth-pollinated niche, and the deliberate design of a hawkmoth-pollinated flower demonstrates a new, predictive method for studying pollination syndrome evolution.

---

Darwin called the dramatic radiation of the *ca*. 250,000 flowering plant species “an abominable mystery,” though he recognized the potential role of the strong coevolutionary relationships between plants and their pollinators (*1*). It is now clear that animal pollination is responsible for high rates of speciation in the flowering plants (*2*). Shifts between pollinator guilds (*e.g*., bumblebees, hummingbirds, hawkmoths, bats) often coincide with plant speciation events (*3*), and each pollinator guild is attracted by a different suite of floral traits (*e.g*., color, pattern, shape, nectar reward) collectively known as a pollination syndrome (*4*). Extensive work has identified pollination syndromes among various plant families (*4*), but the detailed genetics of traits involved in pollinator shift-driven plant speciation remain largely unresolved. Have we learned enough about the genetic basis of the origin of flowering plant species to *engineer* a shift in pollinator guilds? Borrowing from Gould’s metaphor of the “tape of life,” (*5*) can we *anticipate* (rather than recapitulate) evolutionary trajectories, and, instead of replaying the tape of life, run the tape in fast forward?

A recurring pattern of pollinator shifts from hummingbird to hawkmoth pollination has characterized plant speciation in many western North American taxa (*3*,*6*), but in the genus *Mimulus* (monkeyflowers) section *Erythranthe* the evolution of hawkmoth pollination from hummingbird-pollinated ancestors has not occurred. “Hawkmoth flowers” share several characteristics with “hummingbird flowers,” including a large volume of dilute nectar and a long tubular corolla. But most hummingbird flowers are red, hence not easily visible to hawkmoths, whose visual sensitivity does not extend into the longer wavelengths (*7*). Hawkmoth flowers are usually white (or pale) and highly reflective (*6*), adapted for detection by crepuscular and nocturnal hawkmoths.

Our goal is to design and synthesize a new *Mimulus* species, pollinated by hawkmoths and reproductively isolated from its red-flowered, hummingbird-pollinated ancestor, *M. cardinalis*. As a first step, we manipulated two flower color loci in *M. cardinalis* and tested the attractiveness of the resulting four color phenotypes (red, yellow, pink, white) to naïve hawkmoths.

The red color of *M. cardinalis* flowers is produced by the combination of high concentrations of anthocyanin (pink) and carotenoid (yellow) pigments. To selectively eliminate either (or both) floral pigments, we crossed *M. cardinalis* to a white-flowered (bumblebee-pollinated) *M. lewisii* homozygous for a recessive allele (*boo1, 8*) unable to produce anthocyanins, and homozygous for a dominant suppressor of carotenoid pigmentation (*YUP, 9*). *M. cardinalis* is homozygous for the alternative alleles (*BOO1 yup*). We self-pollinated the F_1_ (*BOO1/boo1yup/YUP*) to produce a segregating F_2_ population (*N* = 500), from which we recovered the “ancestral” red phenotype (*BOO1 yup*), and the “derived” yellow (*boo1 yup*), pink (*BOO1 YUP*), and white (*boo1 YUP*) phenotypes (Figure 1A). Segregants of each color (*N* = 3 per color) were matched as closely as possible for corolla size and shape and nectar volume.

**Figure 1.**
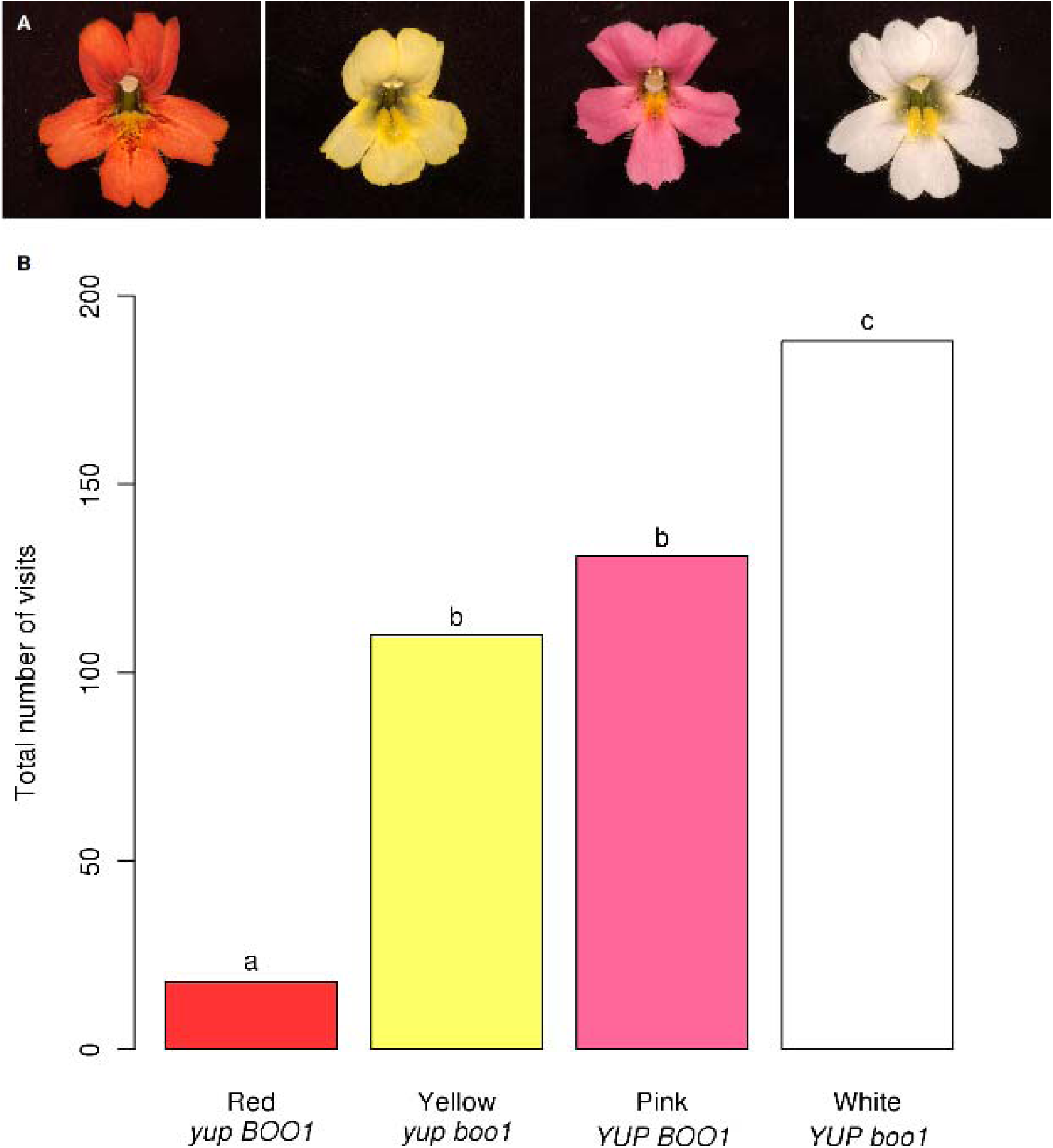
A: Red, yellow, pink, and white *Mimulus* flowers from a single F_2_ population. B: Total number of visits. Letters (a,b,c) indicate significant differences in floral preference red vs. yellow: *X*^2^ = 66.125, *P* = 10^-16^, df = 1; red vs. pink: *X*^2^ = 85.698, *P* = 10^-20^, df = 1; red vs. white: *X*^2^ = 140.2913, *P* = 10^-32^, df = 1; yellow vs. pink: *X*^2^ = 1.8299, *P* = 0.1760, df = 1; yellow vs white: *X*^2^ = 20.4161, *P* = 6.23x10^-6^, df = 1; pink vs. white *X*^2^ = 10.185, *P* = 0.001416).

Using naïve captive-bred female hawkmoths (*Manduca sexta*) in a dimly-lit flight chamber with one flower of each color (Fig. S1), we counted the total number of pollinator visits (see Supporting Material). There were significant differences in visits among the flower color phenotypes (Figure 1B; *N* = 447, *X*^2^ = 134, *P* = 10^-29^). These results indicate that hawkmoths are attracted to flowers with at least one allele substitution step (yellow or pink) from the red flower color characteristic of the ancestral hummingbird-pollinated *M. cardinalis*, and most strongly prefer the two-allele substitution (white).

Testing these four flower color phenotypes with naïve hawkmoths in an experimental chamber has established the remarkably simple genetic basis of phenotypic change required to initiate a pollinator guild shift from hummingbirds to hawkmoths. Observations of pollinator preference and pollen movement in the native environment of *M. cardinalis* using near-isogenic lines for the *YUP* and *boo1* alleles will be needed for a definitive assessment of reproductive isolation between the hummingbird-pollinated ancestral *M. cardinalis* and the rationally designed hawkmoth-pollinated derivatives with yellow, pink, or white flowers.

The classical approach to understanding plant speciation by pollinator shift is *retrospective* – sister taxa with different pollinators are analyzed for differences in key floral traits (*3*,*10*) and their underlying alleles (*8*,*11*) to infer the evolutionary history of divergence from their common ancestor. But perhaps the most stringent test of our understanding of flowering plant diversification is the *prospective* approach we have used here. Darwin famously predicted that the Malagasy star orchid *(Angraecum sesquipedale)*, which has a white flower and *ca*. 35cm nectar spur, must be pollinated by a (then-undiscovered) hawkmoth with a *ca*. 35cm proboscis (*1*). We have shown that critical steps towards the origin of a new, human-designed, hawkmoth-pollinated plant species can, likewise, be predicted based upon a fundamental knowledge of pollination syndromes and genetics.

## Supporting Material

Materials and Methods; References;

**Figure 1:**
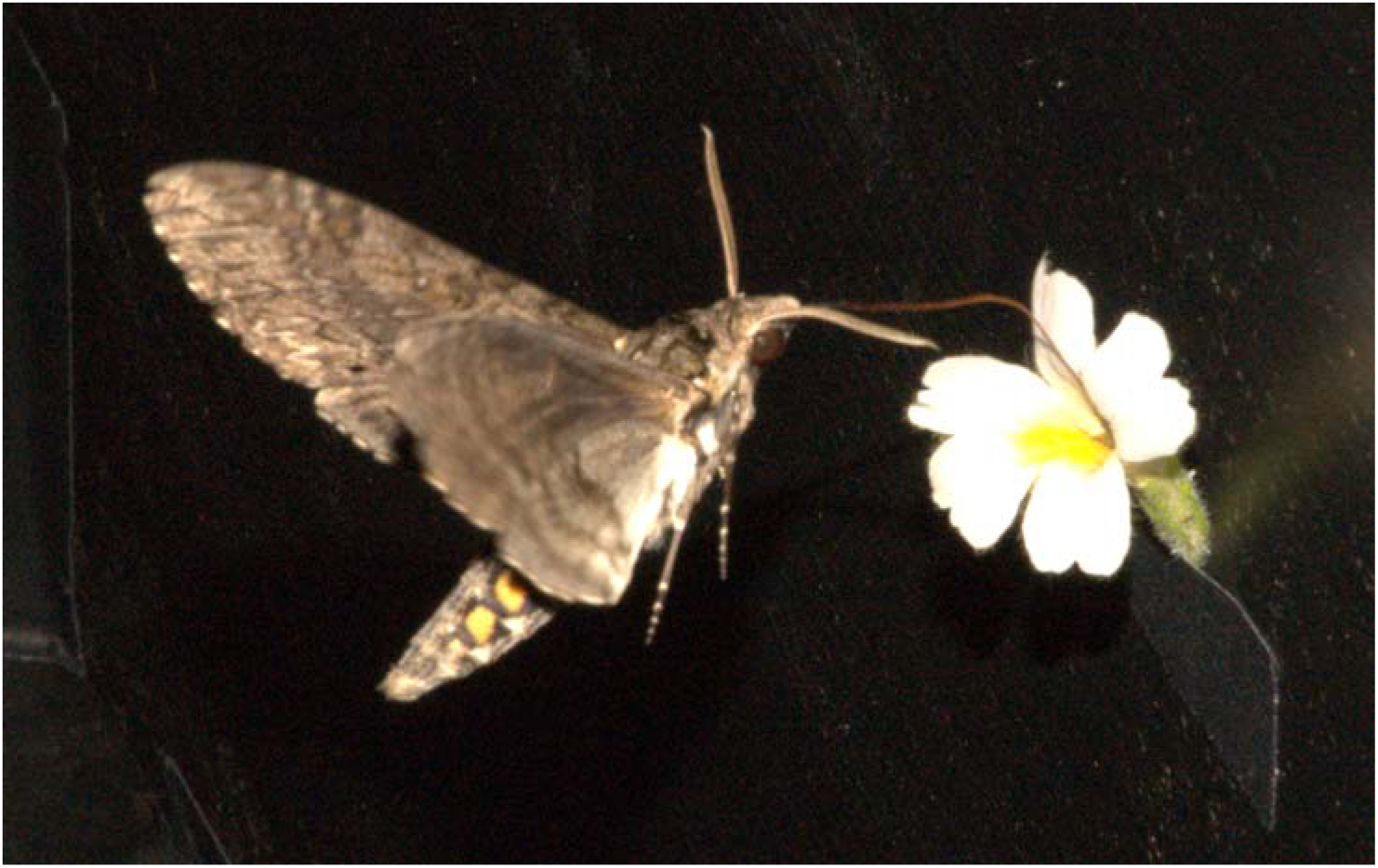
Carolina hawkmoth (*Manduca sexta*) feeding from white *Mimulus* mutant in flight chamber.

## SUPPORTING MATERIAL

### Genetic stocks

*Mimulus cardinalis* Douglas ex Benth. (inbred line CE10, derived by single seed descent from a plant collected along the South Fork of the Tuolumne River, Yosemite, CA) was crossed to *Mimulus lewisii* Pursh (inbred LF10 line derived in the same way from the same area) homozygous for a recessive EMS-induced mutation at the *BOO1* locus, producing an anthocyanin-less flower. A resulting pink-flowered F_1_ offspring was selfed to produce the segregating F_2_ study population. Flowers of four colors were selected, corresponding to the four combinations of alleles at the two flower color loci: red, similar in color to *M. cardinalis* (*yup BOO1*); pink (*YUP BOO1*); yellow (*yup boo1*); and white (*YUP boo1*). Three F_2_ plants of each color were selected based on similarity of flower size, shape (including petal reflexing), and nectar volume.

### Hawkmoth population

Carolina hawkmoths (*Manduca sexta*) were raised on artificial diet (*1*) under controlled conditions at the University of Washington. Hawkmoths were eclosed in full artificial lighting and were not fed or light-cycled prior to the experimental runs. Female hawkmoths eclosed four to six days prior to the experiment (*2*) were used in all experiments.

### Test chamber experiments

Hawkmoths were tested in a 1m x 1m x 70cm chamber constructed of black Coroplast (Coroplast, Dallas, TX) with a clear Plexiglas top for observation. The chamber was located within a darkroom illuminated with red safelight; the chamber itself was illuminated with a single blue-white LED emitting 2 lumens mounted on the Plexiglas top. One flower (including pedicel) of each color was mounted at a height of 50cm on a long side of the symmetrical chamber. Each run was randomized for both flower color at each position and one of three parent plants for each color.

Each hawkmoth was observed until an initial naïve choice (defined as proboscis extension and contact with the floral surface, see *3*) was made. At that point, number of visits until nectar exhaustion or hawkmoth exhaustion were recorded (nectar exhaustion was defined as a visit of one second or less; hawkmoth exhaustion as the hawkmoth becoming unwilling to fly), at which point the hawkmoth was removed and the flowers replaced before a new hawkmoth was introduced. Data were analyzed using a chi-square goodness-of-fit and individual ranking was done with pairwise chi-square tests with a sequential Bonferroni correction (*4*). Visit profiles were examined to rule out initial preference having an effect on further visits (i.e. learning); of 28 moths, 22 visited all three non-red colors, distributed evenly with initial preference ( *X*^2^ = 1.375, *P* = 0.503, df = 2); additionally, 17 of the 28 moths visited another color as often as or more than their initial choice.

